## Supporting Information for "High-throughput fluorescent assay for inhibitor screening of proteases from RNA viruses"

**Supplementary Table 1: The half maximal inhibitory concentration ($IC_{50}$) values determined for inhibitors of the NS2B-NS3$^{pro}$ and PL$^{pro}$.**

| Protease | Inhibitor | $IC_{50}$ (µM) |
|---|---|---|
| NS2B-NS3$^{pro}$ | Aprotinin | 2.4 ± 0.2 |
| | DTNB | 303 ± 54 |
| PL$^{pro}$ | Antabuse | 0.1 ± 0.04 |
| | JB-24 | 1.48 ± 0.39 |
| | 2-MP | 0.82 ± 0.6 |
| | 6-MP | ~ 66 |

**Supplementary Table 2: Amino-acid sequences of substrates for the proteolytic reactions of NS2B-NS3$^{pro}$ and PL$^{pro}$.**

**PL$^{pro}$ substrate: 6xHisTag-mCherry-TEVsite-ISG15-eGFP**

```
MNGSSHHHHHHVSKGEEDNMAIIKEFMRFKVHMEGSVNGHEFEIEGEGEGRPYEGTQTAKLKVTKGG
PLPFAWDILSPQFMYGSKAYVKHPADIPDYLKLSFPEGFKWERVMNFEDGGVVTVTQDSSLQDGEFI
YKVKLRGTNFPSDGPVMQKKTMGWEASSERMYPEDGALKGEIKQRLKLKDGGHYDAEVKTTYKAKKP
VQLPGAYNVNIKLDITSHNEDYTIVEQYERAEGRHSTGGMDELYKSGATENLYFQGAMGWDLTVKML
AGNEFQVSLSSSMSVSELKAQITQKIGVHAFQQRLAVHPSGVALQDRVPLASQGLGPGSTVLLVVDK
CDEPLNILVRNNKGRSSTYEVRLTQTVAHLKQQVSGLEGVQDDLFWLTFEGKPLEDQLPLGEYGLKP
LSTVFMNLRLRGGGTEPGGRSNVSKGEELFTGVVPILVELDGDVNGHKFSVSGEGEGDATYGKLTLK
FICTTGKLPVPWPTLVTTLTYGVQCFSRYPDHMKQHDFFKSAMPEGYVQERTIFFKDDGNYKTRAEV
KFEGDTLVNRIELKGIDFKEDGNILGHKLEYNYNSHNVYIMADKQKNGIKVNFKIRHNIEDGSVQLA
DHYQQNTPIGDGPVLLPDNHYLSTQSALSKDPNEKRDHMVLLEFVTAAGITLGMDELYK*
```

**NS2B-NS3$^{pro}$ substrate: 6xHisTag-eGFP-RSSRRSDLVFS-mCherry**

```
MNGSSHHHHHHVSKGEELFTGVVPILVELDGDVNGHKFSVSGEGEGDATYGKLTLKFICTTGKLPVP
WPTLVTTLTWGVQCFARYPDHMKQHDFFKSAMPEGYVQERTIFFKDDGNYKTRAEVKFEGDTLVNRI
ELKGIDFKEDGNILGHKLEYNAISDNVYITADKQKNGIKANFKIRHNIEDGSVQLADHYQQNTPIGD
GPVLLPDNHYLSTQSKLSKDPNEKRDHMVLLEFVTAAGITLGMDELYKAMGRSSRRSDLVFSVSKGE
EDNMAIIKEFMRFKVHMEGSVNGHEFEIEGEGEGRPYEGTQTAKLKVTKGGPLPFAWDILSPQFMYG
SKAYVKHPADIPDYLKLSFPEGFKWERVMNFEDGGVVTVTQDSSLQDGEFIYKVKLRGTNFPSDGPV
MQKKTMGWEASSERMYPEDGALKGEIKQRLKLKDGGHYDAEVKTTYKAKKPVQLPGAYNVNIKLDIT
SHNEDYTIVEQYERAEGRHSGGMDELYK*
```

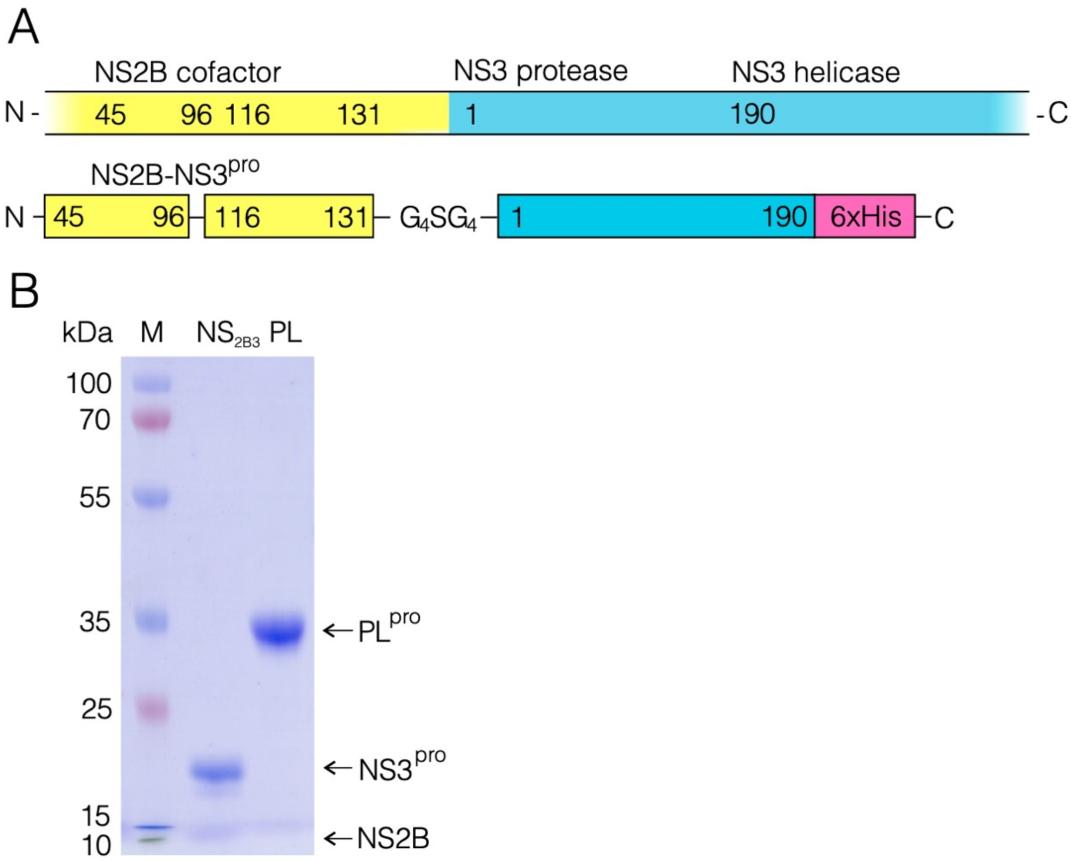


**Supplementary Figure 1: Construct of NS2B-NS3 protease and validation of molecular masses**

(A) To design the NS2B-NS3$^{pro}$, nonstructural proteins NS2B (residues 45-96) and NS3 (residues 116-131) were connected by a flexible glycine-rich linker. A polyhistidine tag was added to the C terminus to enable the purification. (B) Mass spectrum of the NS2B-NS3$^{pro}$. (C) SDS-PAGE gel of the NS2B-NS3$^{pro}$ and PL$^{pro}$ where (M) is a marker, ($NS_{2B3}$) is NS2B-NS3$^{pro}$, (PL) is PL$^{pro}$.

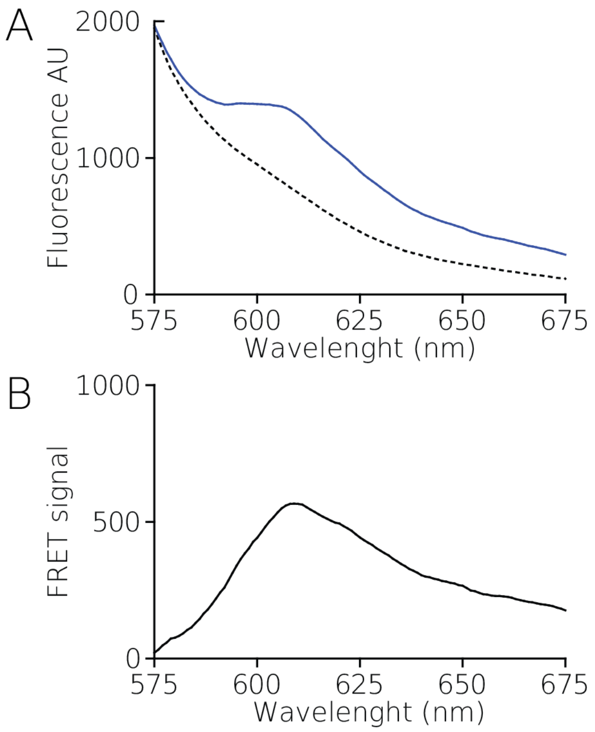


**Supplementary Figure 2: Fluorescent emission spectra demonstrating FRET signal loss for PL$^{pro}$ substrate:**

(A) fluorescence emission spectra of PL$^{pro}$ substrate (solid-blue line) and product (dashed line) measured before and after the reaction with the enzyme (excitation wavelength was 480 nm), (B) FRET emission, the differential spectrum between the intensity of the substrate and product spectra.

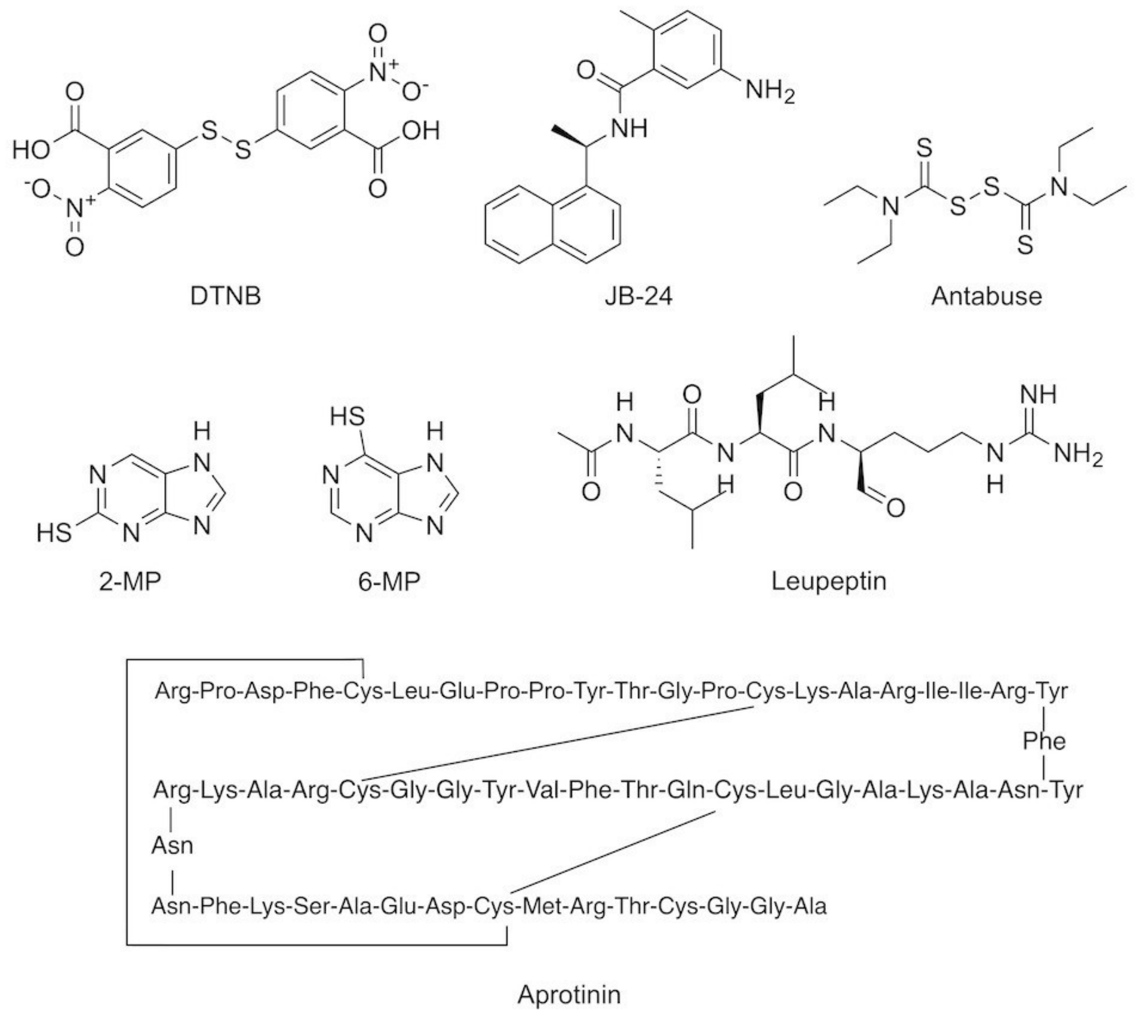


**Supplementary Figure 3: Structures of small inhibitor molecules used in this study**

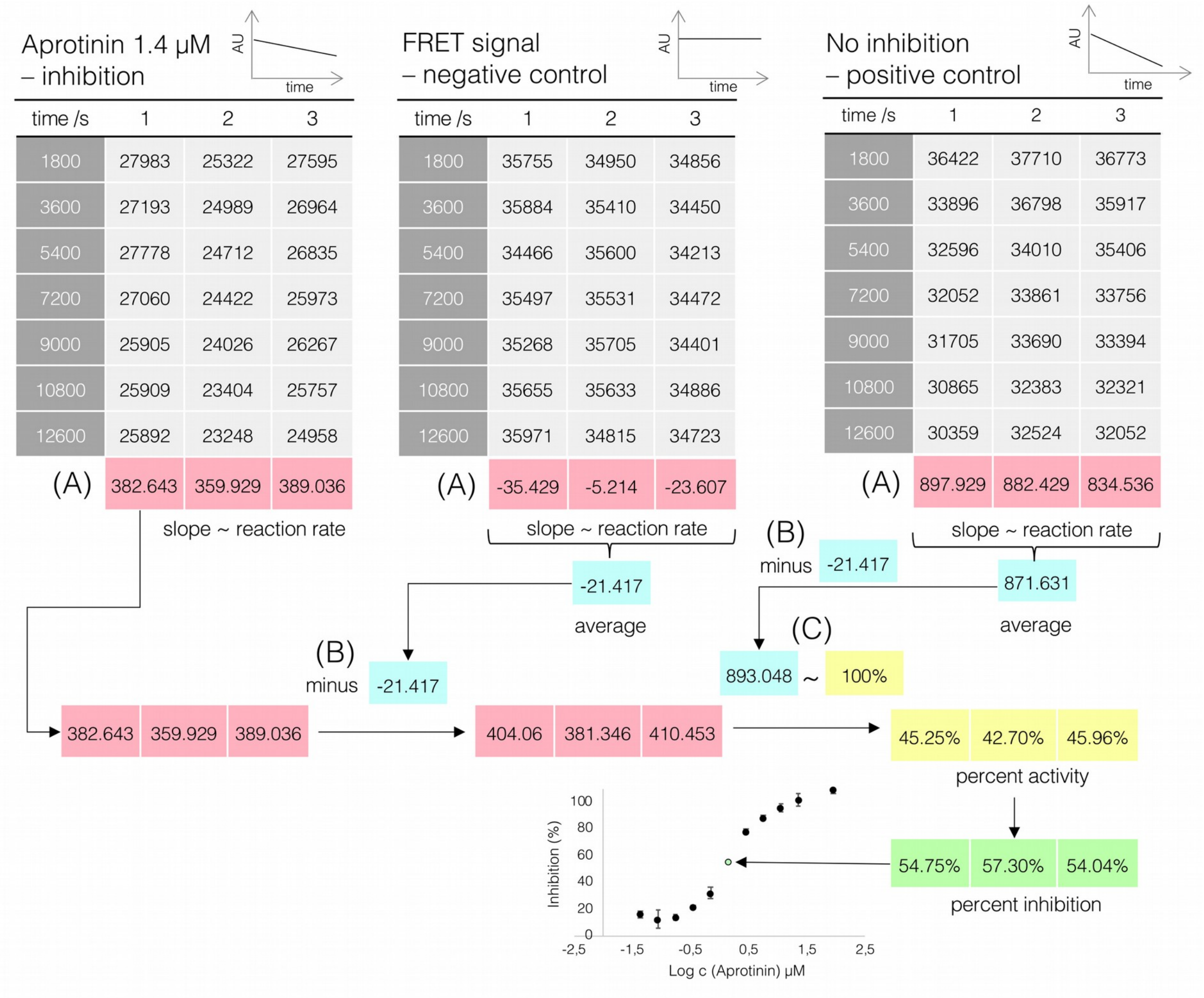


**Supplementary Figure 4: Illustrative evaluation of data for a single set of inhibition reactions (NS2B-NS3$^{pro}$ with Aprotinin as an inhibitor).**

Tables contain illustrative raw data for one experimental set  Each reaction was done in triplicate. (A) The slope of AU versus time was calculated for each well, representing the reaction rate in the corresponding well. (B) Subsequently, the average reaction rate of negative control was subtracted from reaction rates of inhibition reactions and from the average reaction rate of the positive control to correct the data for bleaching and diminishing fluorescence. (C) Corrected rates of inhibition reactions were converted to values of per cent activity and per cent inhibition according to the positive control. The per cent inhibition was plotted against log concentration of the appropriate inhibitor to determine the $IC_{50}$ value of the inhibitor.

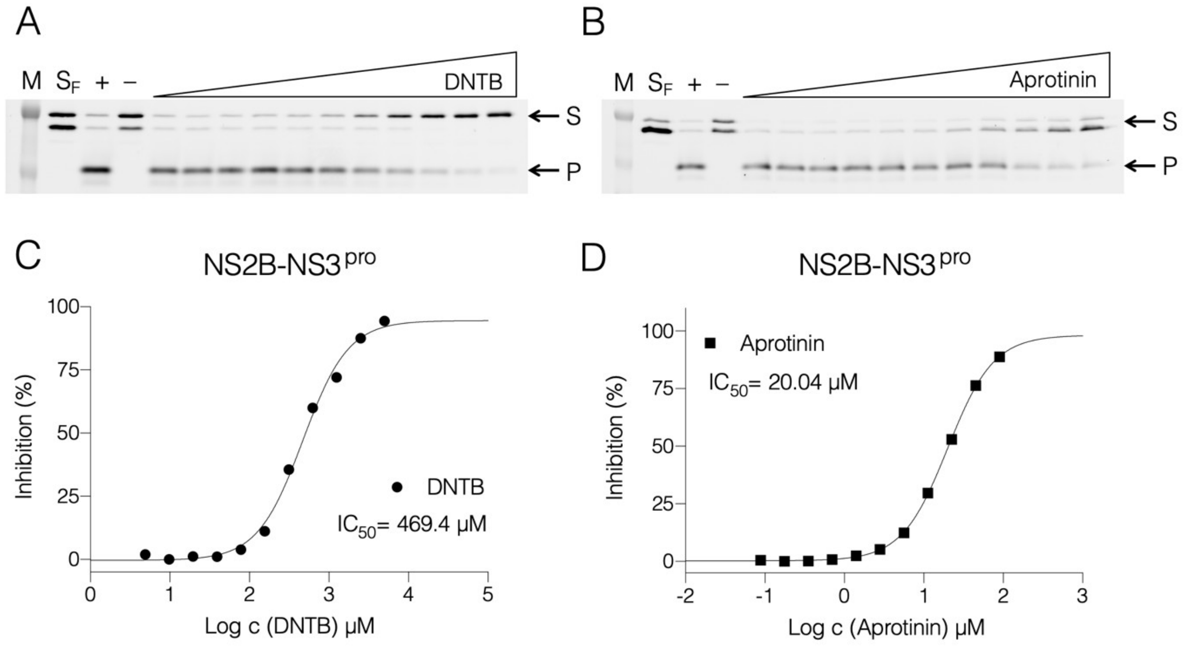


**Supplementary Figure 5: Densitometric analysis of TBEV NS2B-NS3$^{pro}$ inhibition by small molecules resolved on SDS-PAGE.**

Curves derived from densitometric analysis of the SDS-PAGE gels, (A-B) Fluorescence scans of SDS-PAGE gels depicting proteolytic reaction and inhibition by DNTB (A,C) and Aprotinin (D,B), respectively. Substrate conversions were determined, data were normalized to obtain percentage of inhibition and these were plotted against the Log of concentration of inhibitor and fitted with dose-response curve; (M) is a marker, (SF) is fresh substrate at reaction concentration, (+) is positive control reaction where there was no inhibitor present, (–) is negative control reaction where there was no enzyme present, (S) is the substrate, (P) is the product.

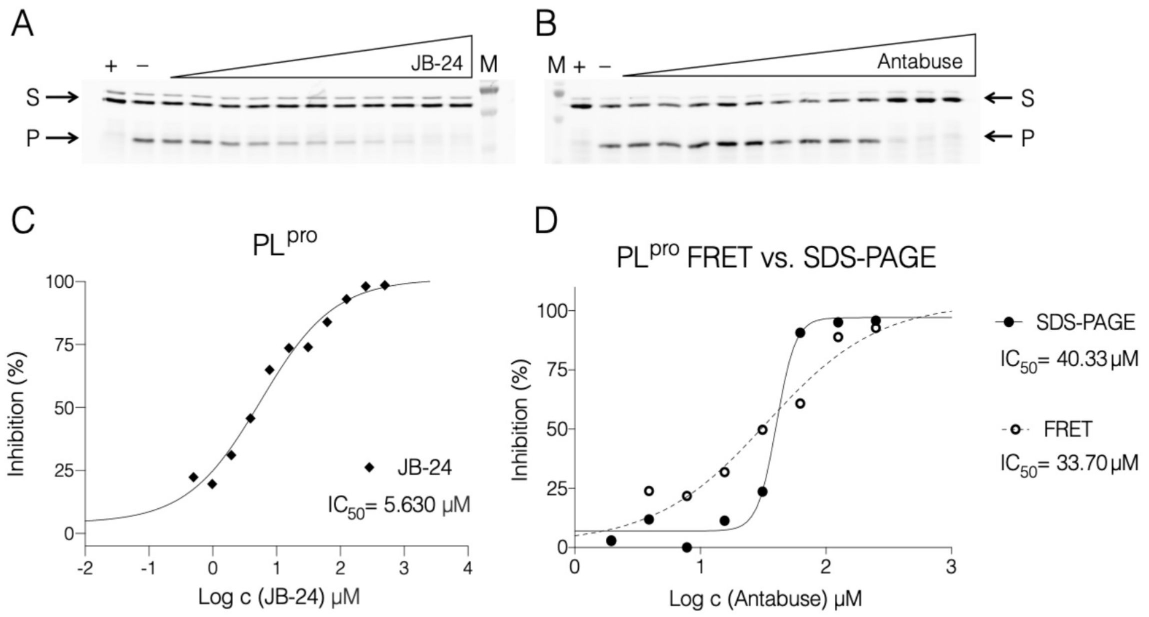


**Supplementary Figure 6: Densitometric analysis of SARS-CoV-2 PLpro inhibition by JB-24 resolved on SDS-PAGE and comparative analysis of FRET-based assay and SDS-PAGE assay with inhibitor Antabuse.**

Curves derived from densitometric analysis of the SDS-PAGE gels, (A-B) Fluorescence scans of SDS-PAGE gels correspond depicting proteolytic reaction and inhibition by JB-24 (A,C) and Antabuse (D,B), respectively. Evaluated enzymatic data from the gels were plotted against log of concentration of inhibitor and fitted to obtain IC50 values; (M) is a marker, (+) is positive control reaction where no inhibitor was present, (–) is negative control reaction where there was no enzyme present, (S) is the substrate, (P) is the product.

(D) The example of a single set of reactions with Antabuse titration assayed by both FRET-based assay and SDS-PAGE gel-based inhibition curve. Data were evaluated as above without averaging. The reaction mixture contained residual reducing agent so it cannot be directly compared with data from Figure 3.

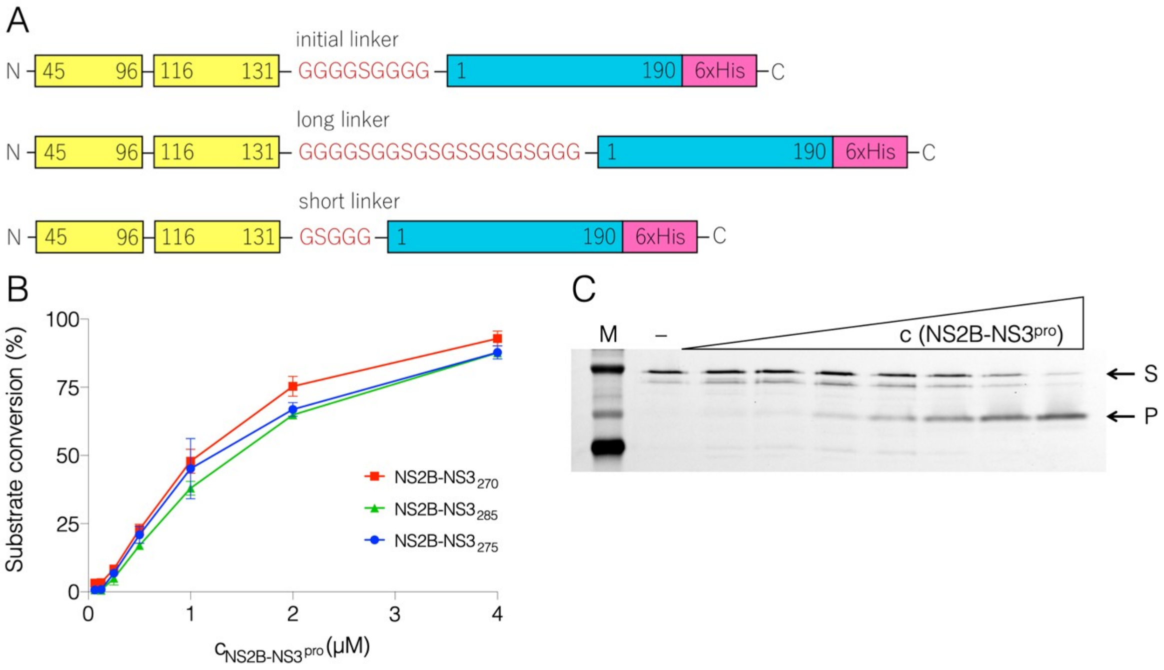


**Supplementary Figure 7: Influence of length of the glycine-rich linker between NS2B and NS3**

(A) Two additional NS2B-NS3$^{pro}$ constructs with different length of the peptide linker between NS2B and NS3 chains were prepared (longer linker $G_4SG_2SGSGS_2GSGSG_3$ and shorter linker $G_1SG_3$). These constructs were used to perform gel based assays with NS2B-NS3 FRET substrates, (B) Graphical representation of substrate conversion with increasing concentration of NS2B-NS3pro with short (red), long (green) or original (blue) length of glycine-rich linker between NS2B and NS3, (C) Fluorescent SDS-PAGE gel from one of the titration of NS2B-NS3$^{pro}$ with the original length of the linker, where (M) is a marker, (–) is a negative control without presence of the enzyme, (S) is the substrate and (P) is the product.
